# BettaAI: a machine-learning object detection model for the quantification of aggressive displays in the Siamese fighting fish *Betta splendens*

**DOI:** 10.64898/2026.09.30.755682

**Authors:** Sara D. Cardoso, Bianca Fusani, Andreia Ramos, Jorge Mendes Maria, David Gonçalves

## Abstract

The Siamese fighting fish (*Betta splendens*) is well-known for its high levels of aggression and complex, stereotyped displays, making it an ideal model for testing deep learning-based tools for high-throughput, unbiased quantification of aggressive behaviour. A deep-learning object recognition model (BettaAI) was developed to detect the species’ aggressive displays, including threat (opercula opening and fin distension), attack (bites) behaviours, as well as surface air breathing, a behaviour correlated with the frequency and intensity of aggression. Videos of males confronting their mirror image were acquired simultaneously with synchronised top and side cameras. Two separate models were trained, with some behaviours identified from top-view frames and others from side-view frames; outputs from both models were combined and post-processed for final quantification. BettaAI performed with high accuracy: F1 scores were 0.99 for the top-view model and 0.93 for the side-view model. Average Precision (AP) at 0.5 IoU exceeded 0.98 for all categories except bites, a rare behaviour in the dataset, which reached 0.865. Comparisons with a manually curated “ground-truth” dataset from videos of 10 males showed strong concordance between automated quantification and manual annotations by expert observers across all behaviours. A case study on 461 males reacting either to a mirror or to a video playback of a conspecific demonstrated the high-throughput and unbiased advantages of BettaAI, with threat displays being more frequent toward the mirror image and attack rates not differing between treatments. This work underscores the potential of BettaAI for automated, bias-free quantification of aggressive displays in *B. splendens*.

## Introduction

Quantification of complex behavioural displays in animals has traditionally relied on manual or semi-manual observation methods (e.g., BORIS, Friard & Gamba, 2016; EthoVision, Noldus et al., 2001). These approaches, while foundational to behavioural research, are time-consuming, error-prone, and subject to inter-observer biases, undermining reproducibility and consistency. Furthermore, the manual nature of these techniques limits the scalability of behavioural studies, making it challenging to process large datasets or detect subtle behavioural patterns that require high temporal or spatial resolution.

Recent advances in machine learning (ML), particularly in deep learning, have provided a powerful alternative to traditional methods of behavioural quantification. Deep learning models excel at identifying and classifying patterns in large datasets, offering scalable, efficient, and objective solutions to behavioural research. These tools have already demonstrated remarkable success in studies across diverse animal taxa (Fazzari et al., 2025). For example, PoseNet (Kendall et al., 2015) and DeepLabCut (Mathis et al., 2018), popular pose-estimation models, have been widely applied to track and analyse locomotion in rodents and zebrafish. In rodents, deep learning has been used to identify grooming, rearing, and social interactions with high precision (Giancardo et al., 2013; Hsieh et al., 2024; Lapp et al., 2023). In insects, ML models have automated the quantification of aggression (lunge) and courtship behaviours (circling, following, attempted copulation, and copulation), in *Drosophila melanogaster* (Yadav et al., 2025). In fish, the utility of ML tools is particularly evident in studies of group dynamics and social behaviours. For instance, Pérez-Escudero et al., (2014) developed idTracker, a software capable of reliably identifying and tracking multiple individual zebrafish in a group, enabling detailed analyses of their interactions and movement patterns. Similarly, Mathis et al., (2018) used DeepLabCut to analyse zebrafish swimming trajectories in three dimensions, revealing subtle variations in movement associated with stress and environmental changes. Deep learning has also been applied to study predator-prey interactions, with models trained to detect and classify events such as attacks and escapes in real-time (Wei et al., 2025). These examples illustrate how ML tools can provide unbiased, high-throughput analyses of complex behaviours that were previously labour-intensive to quantify, standardising results within and across studies and laboratories.

In this study, we developed a YOLO-based deep learning object recognition model, BettaAI, to quantify the aggressive behaviours of male Siamese fighting fish *Betta splendens*. YOLO, a state-of-the-art object detection framework, is well-suited for tracking multiple behaviours in complex environments and is being increasingly applied for the study of animal behaviour (e.g., Hu et al., 2021; Sun et al., 2021). Using synchronised top- and side-view video recordings, we captured three-dimensional information on fish behaviour and position and trained separate models for each perspective to detect opercular opening, fin distension, bites, surface air breathing, and the position of the head and tail. By integrating outputs from these models, we aimed to achieve precise and high-throughput quantification of the species’ aggressive displays.

The Siamese fighting fish has long been used as a model to study aggressive behaviour due to the intrinsically high levels of male aggression. In nature, males establish territories where they build bubble nests to attract females for mating and defend them vigorously against competitors. In Southeast Asia, this high level of aggression has been recognised for centuries, with the species being used for staged paired fights and “winner” strains developed across generations (Ramos & Gonçalves, 2019). During combats, males show stereotyped and visually striking displays, including opercular openings, fin distension, charges, and bites (Ramos & Gonçalves, 2019). These behaviours are easily elicited under controlled experimental conditions, such as exposure to a mirror image or video playback of conspecifics (Alex et al., 2023; Ramos et al., 2021), avoiding the ethical concerns of male-male fights. The predictability and stereotyped nature of aggressive behaviours in *B. splendens* are particularly suitable for developing and validating ML-based tools for behavioural quantification. Combined with recent advances such as the sequencing of the species genome (Zhang et al., 2022) and the possibility of genomic editing (Palmiotti et al., 2023), BettaAI adds to the toolbox available to study the biology of aggressive behaviour in *B. splendens*.

The objectives of this work were threefold: (1) to develop a robust, high-throughput ML model for tracking and quantifying aggressive behaviour in *B. splendens*; (2) to validate the model’s performance against a manually annotated dataset; (3) to apply the model in an experimental context to evaluate its practical applicability. By achieving these aims, BettaAI demonstrates the transformative potential of artificial intelligence in providing unbiased, scalable, and efficient analyses of complex animal behaviours, with particular emphasis on the behavioural diversity of fish.

## Methods and Materials

### Study subjects and housing conditions

All animals used in the experiment were adult *Betta splendens* siblings belonging to the F2 generation derived from an initial cross between a wild-type female and a male from a strain selected for winning fights (for details on these strains, see Ramos & Gonçalves, 2019). The use of F2 hybrid fish for the development of the deep-learning model aimed to capture phenotypic variation associated with the wild-type and fighter parental strains, potentially increasing the generalisation of the model. Fish were reared in mixed-sex groups of approximately 50-100 individuals per tank and housed according to breeding batch. Most fish were subsequently transferred to individual 9W x 9D x 20H cm aquaria between 4 and 8 months post-hatching. Behavioural testing was conducted in adult fish between 14 and 20 months post-hatching, with most individuals tested at 18-20 months. Fish were fed twice a day with newly hatched *Artemia* for the first four weeks post-hatching and were subsequently fed commercial dry food (Golden Pearls and Atison’s Betta Pro) in the morning and live adult *Artemia* in the afternoon, except on the day of the experiment where they were not fed. Conditions were similar under group and individual housing conditions, with water temperature kept at 28 ± 1 °C, the photoperiod set to 12L:12D, and tank water supplied by reverse osmosis (RO) water with salinity (Instant Ocean) kept at 250 ppm and conditioned with Indian almond tree leaves.

### Video recordings

Images used for the training and validation of the model were extracted from video recordings. The setup used to acquire the videos consisted of a tank of 25 W X 12.5 D X 20 H cm filled with a volume of approximately 3.3 L of system water (RO water adjusted to a salinity of 250 ppm using Instant Ocean) and placed inside a recording box diffusely illuminated by LED lights. Each box had one side and one top Raspberry Pi Camera Modules (V2) set to a resolution of 1640 x 922 px and at 30 fps. Each camera was connected to an independent Raspberry Pi 4 Model B, which recorded the video. Trials consisted of a 30-min acclimation period, followed by 30 min of exposure to a mirror placed at one end of the tank to elicit aggressive behaviour. During acclimation, the mirror was concealed behind a smart screen that remained opaque and became transparent when activated, allowing the fish to view its own mirror image without any physical manipulation of the experimental setup. The smart screen was controlled by one of the Raspberry Pi boards via a relay switch (Fig. 1), following the automated stimulus-presentation approach previously used in *B. splendens* behavioural experiments (Alex et al., 2023). Trials were initiated from a remote station that triggered simultaneously the two Raspberry Pi boards allowing synchronised video recording from the two cameras. This also eliminated the need for further manipulation of the experimental setup during the trial, minimising potential disturbance to fish behaviour. After each trial, fish were returned to their home aquaria.

**Fig. 1.**
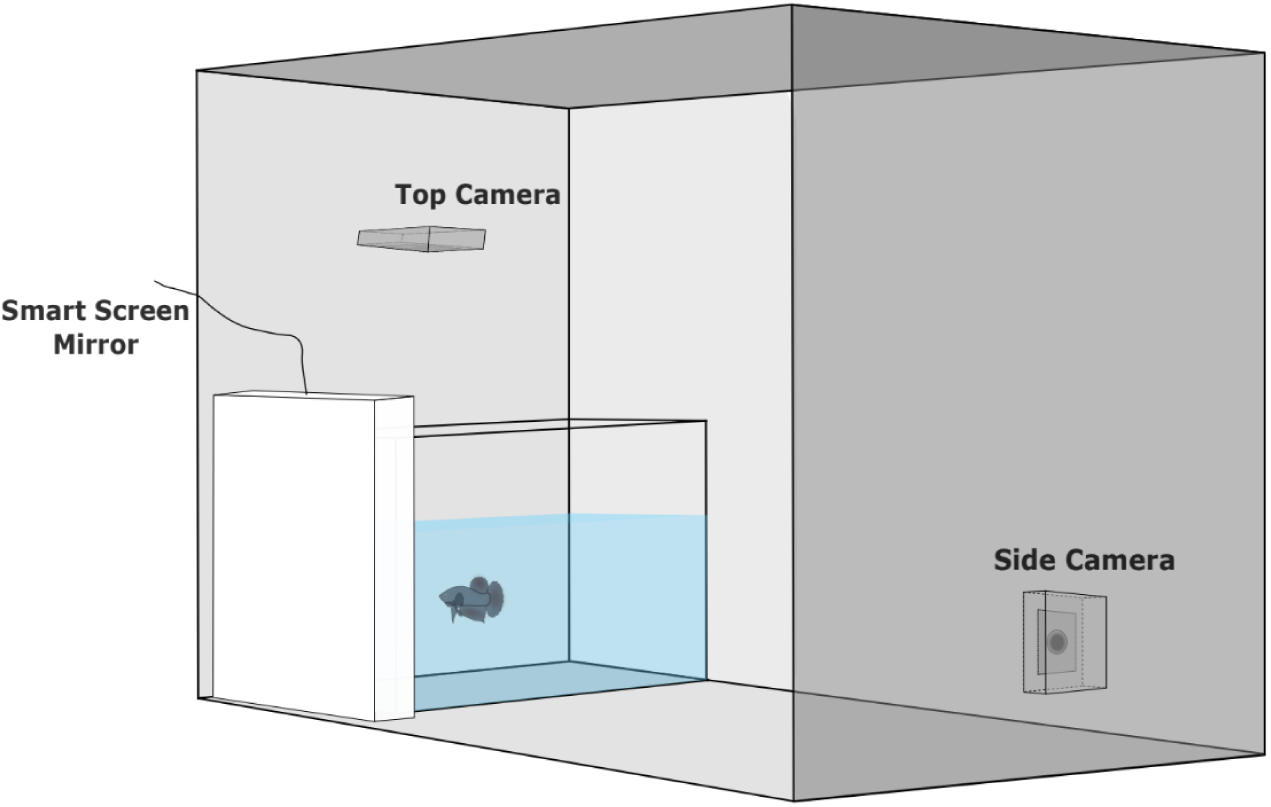
Representation of the experimental setup used for behavioural recording. The focal fish was exposed to a mirror positioned behind a smart screen that remained opaque during acclimation and became transparent at the start of the behavioural trial. The behaviour was recorded simultaneously using top and side cameras. Illumination was provided by a diffuse LED strip.

### Model training

Two separate YOLOv8 nano (Ultralytics v8.1.29; Jocher et al., 2023) object-detection models, pre-trained on the COCO dataset (Lin et al., 2014), were trained using video frames captured by the top and side cameras. Ten object classes were defined across the two models, with six classes per model and two classes shared between models (Fig. 2A). Aggressive behaviours included threat displays such as the “opercula opening” and “distension of the unpaired fins”, through which fish increase their apparent body size to the opponent, and attack behaviours such as “bite” and “charges” (Simpson, 1968). In addition, “air breathing” was also included as a potential indicator of overall metabolic activity (da Silva et al., 2025), which correlates with the frequency of aggressive behaviour (Alton et al., 2012). To determine the three-dimensional position and orientation of the fish, classes were added for different body regions, including “body”, “head”, “center”, and “tail”. Finally, the fish reflection in the aquarium glass walls was also labelled to improve the discrimination between the focal fish and its reflection.

**Fig. 2.**
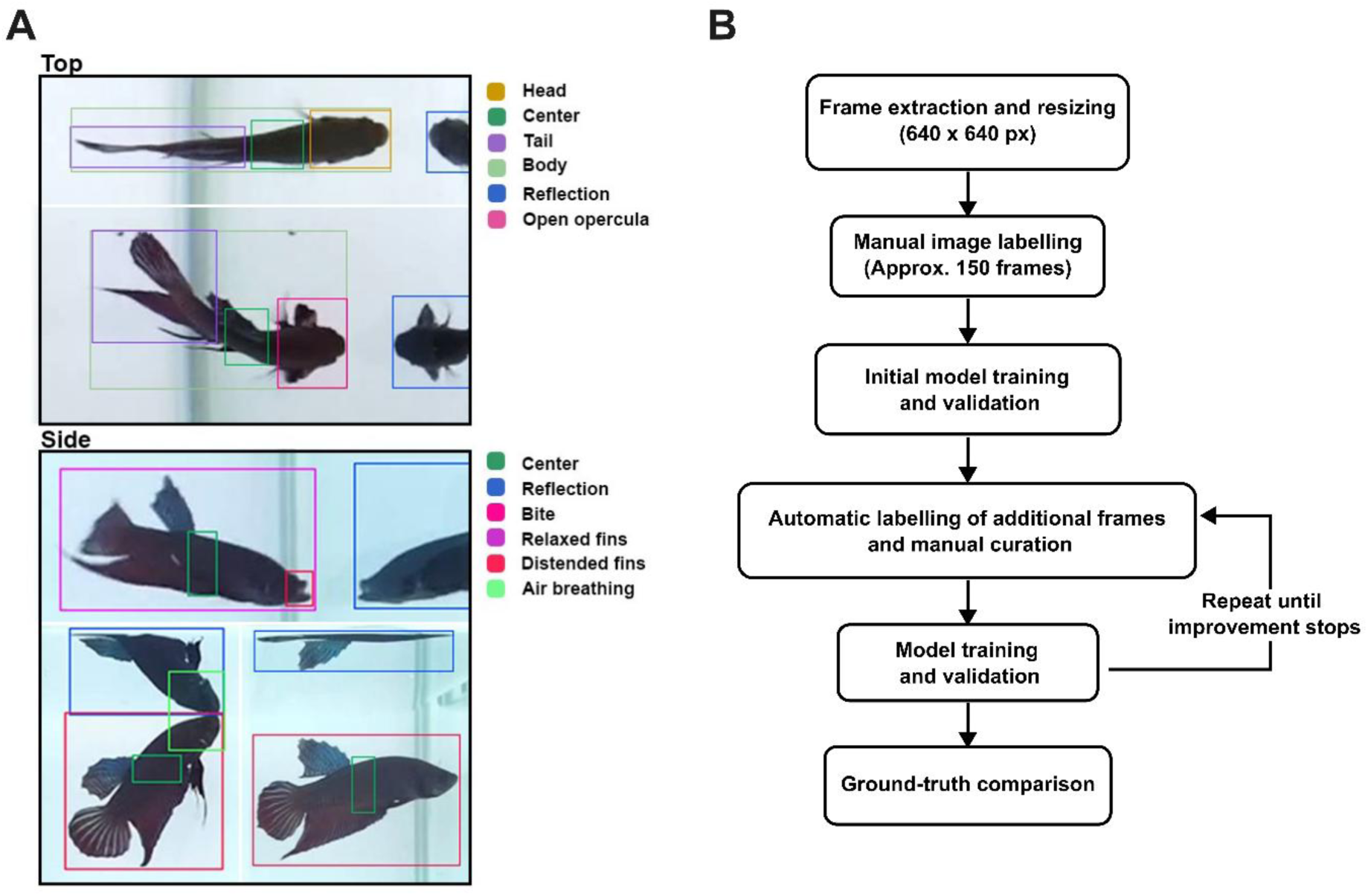
A) Object classes, used to train the top- and side-view models. Each object class is represented by different colour to simplify the visualisation and recognition during the training. B) Flowchart showing the workflow used for training and validation of the model.

The workflow for the two models was the same: (1) frames were randomly sampled from the videos of 13 males for training and 10 different males for validation, so that individuals used for validation were not represented in the training set. Frames were resized to 640 x 640 pixels to standardize input dimensions, without any further preprocessing; (2) a relatively small number of frames (app. 150) were manually annotated using a locally deployed instance of the Computer Vision Annotation Tool (CVAT v2.13.0; CVAT.ai Corporation, 2024) for the different features to be extracted from the top and side images; (3) a first model was trained with this dataset for inference and validated; (4) the initial model was used in a script to automatically generate annotations for additional frames, which were manually reviewed and corrected; (5) this larger dataset was used to train and validate a second model; (6) the previous two steps were repeated until improvement was no longer observable (Fig. 2B). Final datasets comprised 6,920 training and 3,997 validation images for the top-view model, and 8,994 training and 4,002 validation images for the side-view model (Supplementary Table S1). Data augmentation was applied to increase the diversity of the training: mosaic (p = 1.0, disabled for the last 10 epochs), horizontal flipping (p = 0.5), HSV colour-space jitter (hue = 0.015, saturation = 0.7, value = 0.4), translation (0.1) and scaling (0.5), Blur (p = 0.01, blur limit = 3-7), MedianBlur (p = 0.01, blur limit = 3-7), ToGray (p = 0.01), and contrast-limited adaptive histogram equalisation (CLAHE; p = 0.01, clip limit = 1-4, tile grid size = 8 x 8). Vertical flipping was disabled; MixUp, copy-paste and rotation were not used. At each training step, datasets were manually curated to balance the frequency of the different categories, although this was not possible in all cases. For example, while “head” or “tail” were present in all images, “bites” or “air breathing” were infrequent behaviours and had a lower representation (Supplementary Table S1).

Training was conducted on an NVIDIA RTX 3090 GPU using the PyTorch framework. Training performance across epochs was visually assessed with TensorBoard. The final models were trained for 1000 epochs. Model performance was assessed using metrics such as precision, recall, F1 score, and mean Average Precision (mAP), and the best model was selected based on its performance on the validation set.

### Inference and post-processing

For three-dimensional positioning, X and Y coordinates were extracted from the top frame and the Z coordinate from the side frame. The two fixed orthogonal cameras, providing vertical and horizontal views of the aquarium, were calibrated manually by positioning a reference object at multiple known locations within the tank, recording its corresponding image-plane XY coordinates in each camera view, and deriving a calibration factor that was subsequently used to convert pixel coordinates into real-world spatial coordinates. Coordinates for the tank vertices were also extracted, which allowed the three-dimensional calculation of the water volume in the test tank and the relative position of the fish in relation to it. Regions of interest (ROIs) were defined to calculate event timing and frequency across different areas of the test arena. For some variables, thresholds and additional filtering rules were applied before the final results were calculated. For example, in addition to the confidence detection threshold, minimum duration and interval of independent behavioural events were defined. In addition, behaviours such as fin distension are only visible when the fish is oriented laterally to the side camera. Therefore, the angle between the body line and the side camera was extracted from the top frame to identify which of these behaviours could be reliably assessed. Inference was conducted in parallel for the two models, after which data from both models were combined and the post-processing filtering rules applied (Supplementary Table S2). This allowed flexibility in assessing the potential impact of different post-processing rules on the final results, in addition to facilitating quick adjustments to different setup conditions. Post-processed values were used to produce a video overlay of the different behaviours for manual verification of the tracking and behavioural quantification (Supplementary Video 1), allowing fine-tuning of the post-processing settings.

### Ground-truth comparison

A ground-truth dataset was prepared from videos of 10 males that were not used for model training or validation. The same experimental setup previously described was used to record the videos. After a 30-min acclimation period, aggressive behaviours towards the mirror image were quantified during the first 10 min of mirror exposure. Two expert observers independently quantified the following behaviours using Python Video Annotator (Ribeiro et al., 2019): duration and frequency of opercular displays, duration and frequency of unpaired fin distension, frequency of bites, frequency of charges and frequency of air breathing. The values obtained by each observer and by BettaAI were compared to assess agreement and reliability between manually scored and automated behavioural quantification.

### Case-study

The applicability of BettaAI for high-throughput data collection was tested by quantifying the behavioural responses of 503 *B. splendens* males during mirror and video playback tests. Final analyses were restricted to 461 males where complete 10-min recordings for both tests were achieved. These fish belonged to the same F2 cohort used for the model-training and were tested in the previously described mirror-aggression setup. The behaviour was quantified during the first 10 min following stimulus presentation, after the 30-min acclimation period. For the video playback test, the mirror was replaced by an LCD screen positioned at the same end of the focal-fish tank. The video playback setup followed the one described in Alex et al. (2023), with the LCD screen remotely controlled using a Raspberry Pi 4 Model B. During the 30-min acclimation period, a white screen was displayed on the LCD, after which the stimulus video was automatically initiated. Recordings used to construct the video playback stimulus were obtained following the same procedure as described in Alex et al. (2023). In the present study, the final stimulus video consisted of a 30-min composite of 5-min segments from six different males engaged in fights, filmed from the opponent’s perspective. The segments were arranged in temporal sequence to preserve the typical progression of fighting behaviour in this species, and transitions between segments were edited so that fish position within the tank remained as similar as possible. The same stimulus video was used throughout the experiment. Each male was tested in both behavioural tests, and the order of the two tests was balanced across fish, with fish tested within the same week.

### Statistics

Statistical analyses were performed in Python 3.13.15. Inter-observer reliability was first assessed by comparing the behavioural measurements obtained independently by the two expert observers. Agreement and consistency were assessed using intraclass correlation coefficients (ICCs), together with Pearson correlation coefficients and paired-samples t-tests. The measurements obtained by the two observers were subsequently averaged to provide a reference against which the automated measurements generated by the BettaAI were evaluated. Agreement and consistency between the expert observers’ average and BettaAI were again tested using ICCs, Pearson correlation coefficients, and paired-samples t-tests. Agreement was additionally evaluated using Bland-Altman analysis, in which the differences between paired measurements were plotted against their mean. The mean difference was used to estimate systematic bias, and the 95% limits of agreement were calculated as the mean difference ± 1.96 standard deviations of the differences.

Paired-samples t-tests were conducted to assess quantitative differences in the response towards the mirror image and video playback of the conspecific. Pearson correlation matrices for the behavioural responses in each group were calculated, and the Benjamini-Hochberg procedure (Benjamini & Hochberg, 1995) was applied to correct for multiple comparisons among pairwise correlations. Additionally, to determine whether the correlation structure of behaviours differed between the two groups, a Quadratic Assignment Procedure (QAP) correlation test was conducted (Krackhardt, 1988). The observed correlation between the matrices was computed, followed by a QAP permutation test involving 1,000 permutations. It should be noticed that the null hypothesis of the QAP test posits that there is no association between the correlation matrices, with a p-value smaller than 0.05 suggesting they are not significantly associated.

### Ethical note

All methods adhered to the ASAB/ABS (Guidelines for the treatment of animals in behavioural research and teaching», 2012). This study followed the ethical guidelines enforced at the Department of Science and Environment of the University of Saint Joseph and work with the species is approved by the Division of Animal Control and Inspection of the Civic and Municipal Affairs Bureau of Macau, license AL017/DICV/SIS/2016.

## Results

### Model performance against the validation dataset

The two models trained to detect aggressive behaviour in *B. splendens* demonstrated good performance, as evidenced by high precision, recall, and F1 scores across most behavioural classes. For the top-view model, the overall mean Average Precision (mAP) at 0.5 IoU for all classes was 0.993, with Average Precision (AP) values ranging from 0.991 for “Open opercula” to 0.995 for “Tail”, “Center” and “Body”. The normalised confusion matrices showed almost perfect classification for all object classes (Fig. 3A), with the lowest value for the opercula opening, which was classified as head in 4% of cases. The distinction of when an operculum is considered open, or the head otherwise, in a single frame is intrinsically difficult as it depends on the angle of opercula opening and position of the head (Fig. 3B).

**Fig. 3.**
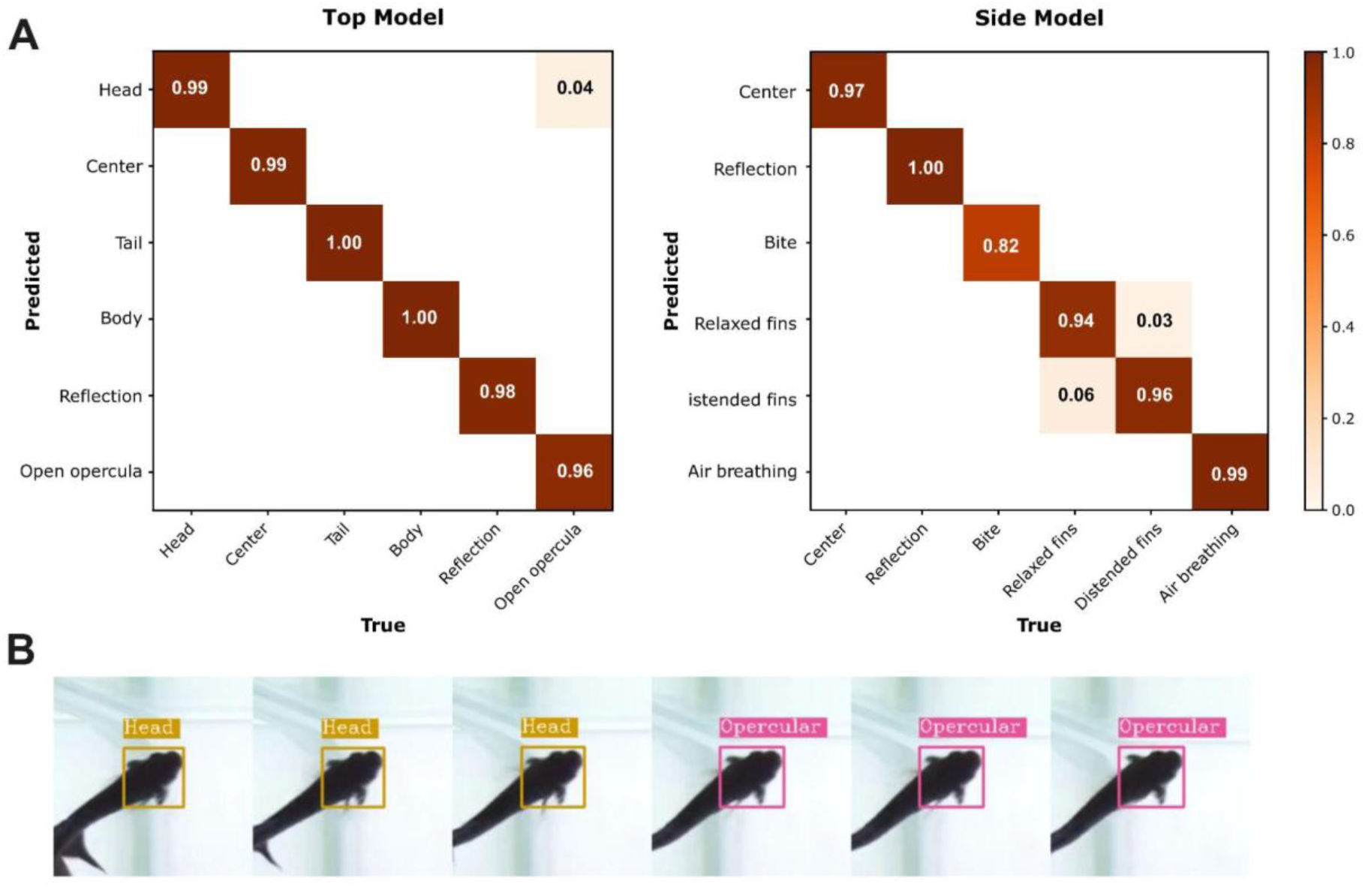
A) Normalised confusion matrices for the top-view (left) and side-view (right) models; B) six sequential frames showing the detection of the head or of open opercula.

A misclassification of 4% is considered low, in particular because post-processing rules (Supplementary Table S2) filtered out most of the misclassification cases. The F1-confidence curve demonstrated an F1 score of 0.99 at the optimal confidence threshold of 0.322, for all classes combined. The training loss for box regression started at 1.0 and converged to 0.4, while the validation box loss started at 0.8 and converged to 0.3. The training classification loss began at 1.5 and decreased to 0.5, with the validation classification loss showing a similar trend from 0.8 to 0.2. The distribution focal loss followed a decreasing trend, indicating improved localisation accuracy over time.

For the side-view model, mAP at 0.5 IoU for all classes was 0.964, indicating slightly lower performance than for the top-view model. This was mainly because of a lower AP for “Bite” of 0.865, with AP values of the remaining categories varying from 0.978 for “Distended fins” to 0.995 for “Reflection”. “Bite” was a rare event in the dataset (Supplementary Table S1), which likely explains the lower accuracy for the detection of this behaviour. An F1 score of 0.93 at the optimal confidence threshold of 0.342 was obtained for all classes combined for the side-view model.

The training loss for box regression started at 1.2 and converged to 0.4, while the validation box loss started at 0.8 and converged to 0.3. Similarly, the training classification loss began at 2.0 and decreased to 0.5, with the validation classification loss showing a similar trend from 0.8 to 0.2. The distribution focal loss also followed a decreasing trend, indicating improved localisation accuracy over time.

### Model performance against the ground truth dataset

Results for the ground truth dataset were first compared between manual annotations by the two expert observers. Agreement was highest for the frequency of air breathing (perfect concordance) and lowest for the frequency of distension of the fins (ICC agreement score = 0.72; ICC consistency score = 0.73). Across all behaviours, the F-tests confirmed the significance of the ICC results (p < 0.01), demonstrating strong inter-observer agreement and consistency. Pearson’s correlation coefficient was above 0.93 for all variables. However, paired t-tests were significant for the frequency (p = 0.031) and duration (p = 0.020) of distension of the fins and close to significant for the time with open opercula (p = 0.058), showing a systematic difference in the classification of these behaviours by the two observers (Supplementary Table S3). The results from the two expert observers were averaged for comparison against BettaAI. Agreement and consistency scores were high between the average of the two expert observers and BettaAI. ICC scores for agreement and for consistency varied between a maximum of 0.99 for the frequency and duration of opercular displays, to a minimum of 0.81 for the duration of distension of the fins. There was a strong correlation between the human and BettaAI classifications, with correlation scores all above 0.93, with the exception of the duration of distension of the fins, which had a correlation of 0.85. No systematic differences were found between the two classifications, as assessed by paired t-tests (Fig. 4, Table 1).

**Fig. 4.**
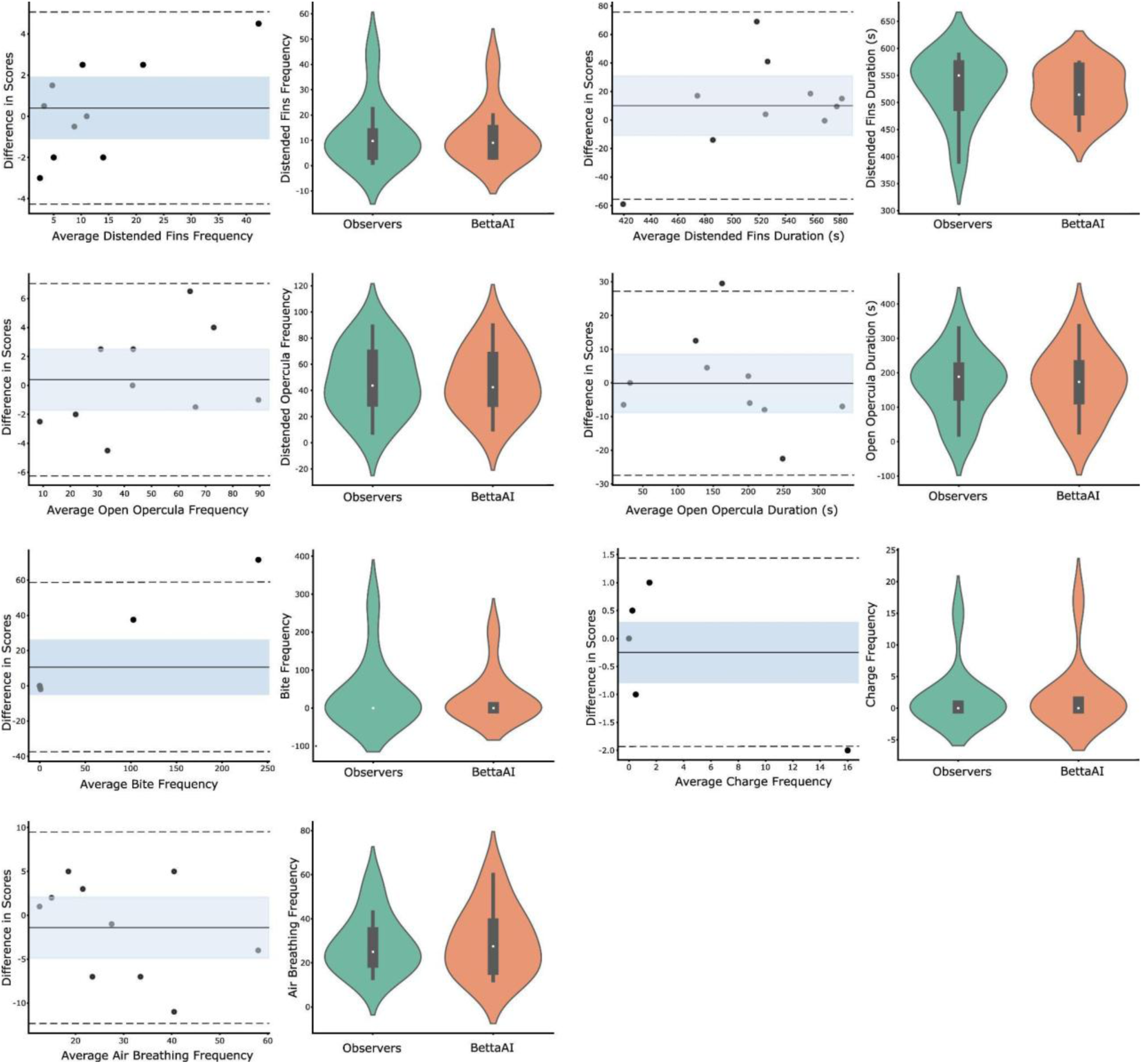
Bland-Altman and violin plots comparing behavioural quantification by the two expert observers’ average (in green) with BettaAI (in orange) for behaviours displayed by 10 male *B. splendens* towards their mirror image during 10-min observation period. In the Bland-Altman plots, the solid horizontal line represents the mean difference between measurements, the dashed lines represent the 95% limits of agreement (mean difference ± 1.96 SD), and the shaded area represents the 95% confidence interval around the mean difference.

**Table 1.** Inter-observer agreement and consistency metrics between the average classification by the two expert observers and BettaAI across the different aggressive behaviours recorded from 10 male *B. splendens* towards their mirror image for 10 min.

|  | ICC agreement | ICC consistency | Pearson correlation coefficient | Paired t-test (d.f. = 9) |
| --- | --- | --- | --- | --- |
| Open opercula duration | 0.990<sup>***</sup> | 0.990<sup>***</sup> | 0.990<sup>***</sup> | -0.022 |
| Open opercula frequency | 0.991<sup>***</sup> | 0.991<sup>***</sup> | 0.992<sup>***</sup> | 0.542 |
| Distended fins duration | 0.813<sup>***</sup> | 0.813<sup>**</sup> | 0.846<sup>**</sup> | 0.936 |
| Distended fins frequency | 0.978<sup>***</sup> | 0.978<sup>***</sup> | 0.987<sup>***</sup> | 0.857 |
| Charges frequency | 0.984<sup>***</sup> | 0.984<sup>***</sup> | 0.990<sup>***</sup> | -0.688 |
| Bites frequency | 0.948<sup>***</sup> | 0.948<sup>***</sup> | 0.999<sup>***</sup> | 1.370 |
| Air breathing frequency | 0.927<sup>***</sup> | 0.927<sup>***</sup> | 0.937<sup>***</sup> | -0.793 |

\* $p < 0.05$ ; \*\* $p < 0.01$ ; \*\*\* $p < 0.001$ .

### Case-study

To illustrate the potential for high-throughput and unbiased estimates of BettaAI, the reaction to a mirror image and to a video playback of an opponent was compared for 461 males. The mirror image elicited more intense threat displays than the video playback footage, as shown by a higher frequency and duration of opercula opening towards the mirror image (paired t-test, open opercula frequency t_(460)_ = 13.03, p < 0.001; open opercula duration t_(460)_ = 13.88, p < 0.001; Fig. 5A). This was also the case for the duration of fin distension (t_(460)_ = 14.46, p < 0.001), with an opposite result for the frequency of this behaviour (t_(460)_ = −4.38, p < 0.001), illustrating the fact that fish spent most of the time with the unpaired fins distended towards the mirror, with shorter and more frequent openings towards the video playback (Fig. 5B).

**Fig. 5.**
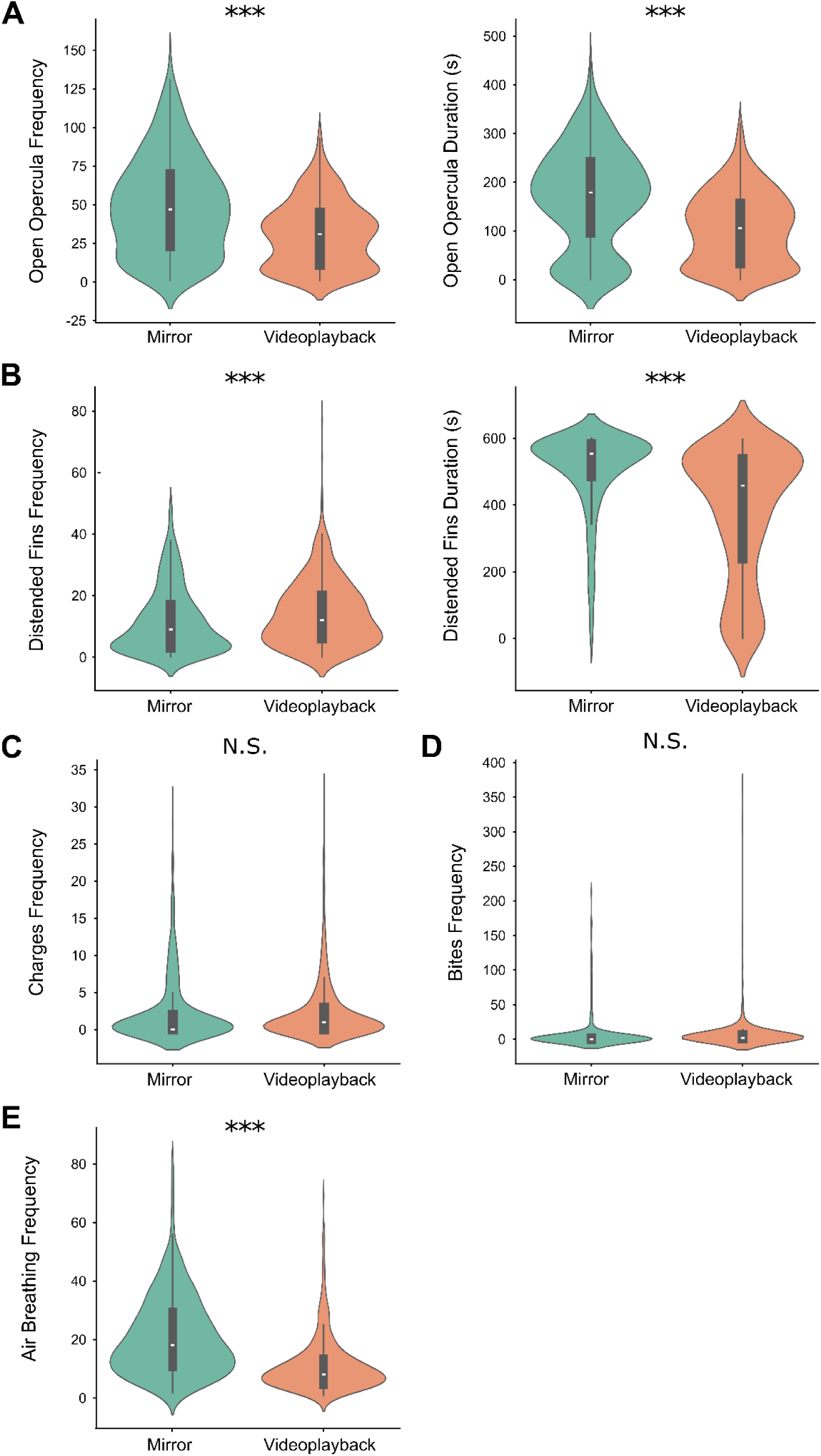
Violin plots for the responses of the same *B. splendens* males (N = 461) to their mirror image (in green) or to a video playback of an opponent (in orange). *** p < 0.001, N.S. p > 0.05.

Interestingly, no differences were recorded for attack behaviours, namely for the frequency of charges (t_(460)_ = 0.86, p = 0.393, Fig. 5C) and of bites (t_(460)_ = −1.20, p = 0.232, Fig. 5D). The frequency of air breathing, which has been shown to be a reliable indicator of metabolic activity in this species (Alton et al., 2012), was higher towards the mirror as compared to the video playback (Fig. 5E) and correlated with most measures of aggression. Overall, there were no differences in the mirror and video playback correlation networks (QAP, r = 0.867, p < 0.001; Fig. 6).

**Fig. 6.**
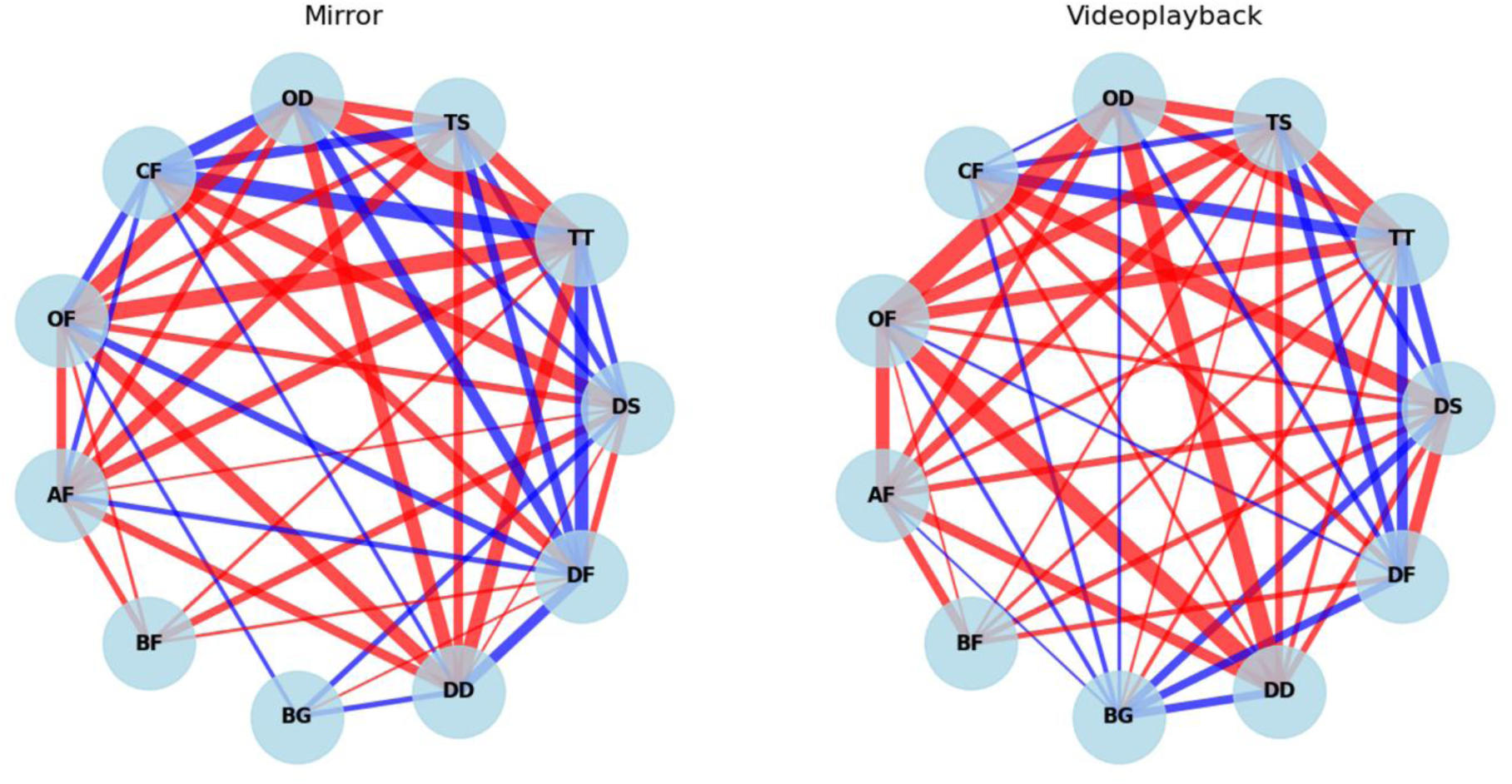
Correlation networks for the different variables quantified by BettaAI for 461 males responding to their mirror image (left) and a video playback of a conspecific (right). OD, Open opercula duration (s); TS, Time at the surface (s); TT, Time near stimulus (s); DS, Distance travelled (cm); DF, Distended fins frequency; DD, Distended fins duration (s); BG, Average body greyscale; BF, Bites frequency; AF, Air breathing frequency; OF, Open opercula frequency; CF, Charges frequency. Positive correlations are indicated with red lines and negative correlations with blue lines, with line thickness proportional to the correlation value. Non-significant correlations are not shown.

## Discussion

This paper describes the development of BettaAI, a machine-learning object detection tool based on two models for quantifying aggressive behaviour in *B. splendens*, and validates its performance against two distinct datasets. The application of the software for analysing multiple videos in high-throughput experiments was also demonstrated. BettaAI achieved robust performance across two different validation approaches. When evaluated against the validation dataset, the normalised confusion matrices for both side- and top-view models showed good performance with low rates of false positives and negatives. The top-view model achieved a mAP of 0.99, with near-perfect classification across all classes in the normalised confusion matrix. The lowest classification accuracy was observed between the “open opercula” and “head” classes, with 4% of “open opercula” instances misclassified as “head”. This is likely because determining when an operculum is open is inherently challenging, as it depends on the angle of both the fish and the opercula relative to the camera. Nevertheless, the post-processing rules implemented improved discrimination between these categories, and a misclassification rate of only 4% is considered low. For the side-view model, the mAP was 0.964, with the lowest AP observed for the “bite” class (0.865). This is likely because biting was a rare event in the training dataset, making it more challenging for the model to learn. The remaining classes all showed high AP values, ranging from 0.978 for “distended fins” to 0.995 for “reflection”. The lowest classification accuracy in the confusion matrix was between “distended fins” and “relaxed fins”, probably because the gradation in fin opening makes labelling challenging.

When compared against the ground truth dataset, where classification was evaluated against two epxert observers, BettaAI again performed well. Agreement and consistency scores between the average of the two expert observers and BettaAI were high across all behaviours, with ICC ranging from 0.99 for opercular displays to 0.81 for the duration of fin distension. Strong correlations were also observed, with Pearson’s coefficients above 0.93 for all variables, except for the duration of fin distension, which had a correlation of 0.85, a value that remains high. Importantly, no systematic differences were found between the human and BettaAI classifications, as indicated by non-significant paired t-tests. This contrasts with the comparison between the two expert observers alone, where significant biases emerged for the frequency and duration of fin distension, and a near-significant bias for time with open opercula, further highlighting how an objective detection model can help standardise recognition of behaviours that are subject to interpretation. These two validation approaches demonstrate that an AI model can reliably recognise and measure aggressive behaviours in fish. One potential refinement concerns the classification of “open opercula” versus “head”, and of “distended fins” versus “relaxed fins”. Although misclassifications between these categories were minimal, in the validation dataset and did not compromise overall model performance, incorporating additional post-processing rules could further enhance precision. For example, the angle between the opercula and the head could be used to determine when the opercula are open (e.g., ≥90°), and a minimum degree of fin distention could be defined for classification as “distended”. These adjustments could further improve the precision of the model, although they are not strictly necessary given its already robust performance.

BettaAI offers several advantages over traditional manual annotation. First, it is much faster to quantify behaviour using the automated process, allowing high-throughput experiments with a significantly higher number of subjects. This was exemplified by an experiment comparing the aggressive response of the same animals to video playback and a mirror image, using a dataset of 461 males. A previous study comparing responses to live and video-playback opponents was limited by manual annotation constraints to 8-10 animals per group (Alex et al., 2023). The comparison performed in that experiment showed differences in the time period of the fight (first or second half) that were independent of stimulus type (Alex et al., 2023). In contrast, the present study reveals that even if the two types of stimuli still enhance aggressive behaviour in bettas, there are subtle differences in threat displays: fish presented with a mirror showed increased opercular opening (both in timing and frequency) and fewer, but longer, distended fin events compared to video playback. These differences, not observed previously, highlight BettaAI’s capacity to detect subtle behavioural patterns through increased statistical power. Furthermore, this enabled the development of stable, stimulus-dependent correlation networks, contributing to a more precise delineation of behavioural responses. Then, as already mentioned, BettaAI allows unbiased and comparable estimates of behavioural results within and across experiments. The two expert observers who manually quantified the ground truth dataset were experienced in behavioural analysis of *B. splendens* aggressive displays, and inter-observer agreement and consistency were generally high. However, a statistically significant bias emerged in the classification of two behaviours, with scores consistently higher for one observer. This finding underscores the need for standardised criteria and procedures to obtain comparable results across studies, a need that automated methods are particularly well-suited to address.

Regarding the study species, *B. splendens* comprises several phenotypically distinct varieties, usually distinguished between long-tails such as “crowntail”, “half-moon” or “veiltail” and short tail (Zhang et al., 2022). *B. splendens*, which is endemic to parts of Southeast Asia, is known for its intrinsic aggressiveness, character that has been recognized for centuries by locals, who began using males in staged fights as a national pastime (Smith, 1945). Winner lines are bred while loser lines are discarded, leading to the establishment of a “fighter strain”. Previous work from the laboratory has investigated the impact of this selection process by comparing wild-type with fighter strains (e.g. Ramos & Gonçalves, 2019, 2022). Fish from these strains have short fins and thus belong to the commonly denominated “plakat” variety. Importantly, the objective of the present study was to develop an automated detection process specifically adapted to the experimental setup and the fish strains routinely used in the laboratory, rather than to create a general model applicable across all Betta varieties. However, the potential application of BettaAI to other strains cannot be excluded without further testing, if at least the experimental setup is maintained in the same way (two cameras). At the same time, if direct application to other strains proves unfeasible, the framework developed here can still serve as a useful and reliable strategy for developing similar tools tailored to different strains in the future. Indeed, the most time-consuming step in creating the model is the labelling phase: as the framework shows (Fig. 2B), a large number of images is required to train the classifier, but time is the only real limitation, as the process itself does not involve any conceptual or technical difficulty. In order to extend the application of the BettaAI model to other contexts, some adaptations would be required depending on the experimental setup, e.g. when using only one camera, working with multiple animals or studying other strains. While some variables quantified by BettaAI are likely unaffected by morphological variation (e.g., position variables, opercular opening, charges, bites), others, particularly fin distension, may require retraining or new training with images of different fish types. Nevertheless, following the procedure outlined here, it should be relatively straightforward to train or retrain the model for other specific Betta varieties.

In conclusion, the BettaAI model proves to be a reliable and efficient tool for the quantification of aggressive behaviour in *B. splendens* when applied in a dual-camera setup (top and side views). By automating the detection and classification of threat displays, BettaAI provides a solid foundation for the broader application of machine-learning approaches in this species. Moreover, the framework developed here can readily be adapted or extended to quantify additional behavioural dimensions, offering a versatile platform for future studies on *Betta* behaviour and potentially other fish species. Although further technical refinements may enhance its precision, the current version already represents a substantial advancement for high-throughput and reproducible behavioural analysis.

## Open Practices Statement

The data and materials, including BettaAI software, source code, trained models, and documentation for all experiments, are available from the corresponding author upon request, and none of the experiments was preregistered.

## Credits

SDC and DG designed the experiments and performed data analysis, and SDC ran the behavioural trials and performed data validation. AR was responsible for data validation. JM developed the software. BF contributed to data visualisation. DG was responsible for supervision, project administration and funding acquisition. All authors contributed to the writing and the revision of the manuscript.

## Acknowledgement

AR had support from the postdoctoral fellowship 0010/2024/STT from the Macao Science and Technology Development Fund (FDCT), SDC from the postdoctoral fellowship 0001/2021/APD from FDCT, and BF from the postdoctoral fellowship 0006/2024/AFJ from FDCT. The study was supported by FDCT project 0088/2023/RIA2.

## Supplementary Materials

**Table S1.** Number of animals, images and labels used for the final training of the top- and side-view models. Images for the two models were extracted from videos of 13 male *B. splendens* for training and 10 male *B. splendens* for validation.

|  | Training | Validation |
| --- | --- | --- |
| Top-view model |  |  |
| Number of images | 6920 | 3997 |
| Head labels | 5001 | 3204 |
| Center labels | 6920 | 3997 |
| Tail labels | 6918 | 3996 |
| Body labels | 6920 | 3997 |
| Reflection labels | 3718 | 2163 |
| Open opercula labels | 1920 | 793 |
| Side-view model |  |  |
| Number of images | 8994 | 4002 |
| Center labels | 8000 | 3989 |
| Reflection labels | 9904 | 4849 |
| Bites labels | 787 | 28 |
| Relaxed fins labels | 3243 | 1502 |
| Distended fins labels | 3636 | 2052 |
| Air breathing labels | 2245 | 552 |

**Table S2.** Variables quantified with BettaAI and the post-processing settings used in this study. For each variable, it is indicated whether information from the top-view model, side-view model, or both models was used.

|  | Top-view model | Side-view model | Post-processing settings |
| --- | --- | --- | --- |
| Aggression |  |  |  |
| Opercular opening | X |  | Minimum duration 3 frames; Minimum interval 30 frames; Minimum confidence 0.85. |
| Fins distension |  | X | Minimum duration 10 frames; Minimum interval 10 frames; Minimum confidence 0.9. |
| Bites |  | X | Minimum duration 1 frame; Minimum interval 1 frame; Minimum confidence 0.7. |
| Charges | X | X | Minimum duration 15 frames; Minimum interval 15 frames; Only when performed in the direction of the stimulus; Only when speed > 0.2 cm/frame. |
| Activity |  |  |  |
| Air breathing |  | X | Minimum duration 3 frames; Minimum interval 30 frames; Minimum confidence 0.7; Only if at the surface. |
| Distance traveled | X | X | Calculated from XYZ position after manual calibration of top and side cameras. |
| Close to stimulus | X | X | Body center in a region of interest of 4 cm close to the stimulus. |
| Away from stimulus | X | X | Body center in a region of interest of 4 cm close to the side opposite to stimulus. |
| At the surface | X | X | Body center in a region of interest of 4 cm close to the surface. |
| At the bottom | X | X | Body center in a region of interest of 4 cm close to the bottom. |
| Orientation | X |  | Line between head/opercula center and body center; Facing stimuli (45-135°). |
| Freezing | X | X | Minimum duration 150 frames; Maximum distance traveled between frames 0.25 cm. |
| Body color |  | X | Grayscale value from the body; side body angle < 10°, extrapolated for other frames. |

**Table S3.** Inter-observer agreement and consistency metrics for comparisons between the classifications of the two expert observers across the different aggressive behaviours recorded from 10 male *B. splendens*. Concordance was perfect for air-breathing frequency.

|  | ICC agreement | ICC consistency | Pearson correlation coefficient | Paired t-test (d.f. = 9) |
| --- | --- | --- | --- | --- |
| Open opercula duration | 0.981<sup>***</sup> | 0.981<sup>***</sup> | 0.988<sup>***</sup> | 2.174 |
| Open opercula frequency | 0.996<sup>***</sup> | 0.996<sup>***</sup> | 0.998<sup>***</sup> | 1.149 |
| Distended fins duration | 0.844<sup>***</sup> | 0.850<sup>***</sup> | 0.934<sup>***</sup> | 2.809* |
| Distended fins frequency | 0.727<sup>**</sup> | 0.740<sup>**</sup> | 0.959<sup>***</sup> | 2.551* |
| Charges frequency | 0.998<sup>***</sup> | 0.998<sup>***</sup> | 0.998<sup>***</sup> | 1.000 |
| Bites frequency | 0.992<sup>***</sup> | 0.992<sup>***</sup> | 0.999<sup>***</sup> | 1.499 |
| Air breathing frequency | 1 | 1 | 1 | NA |

\* $p < 0.05$ ; \*\* $p < 0.01$ ; \*\*\* $p < 0.001$ .

Supplementary Video 1 - Example of top- and side-view recordings after model inference and post-processing. The video illustrates the automated tracking of the fish and detection of behavioural categories. The top-view shows body orientation, together with the detection and labelling of specific behaviours (e.g., Open opercula), while the side-view provides complementary detection of behaviours visible from the lateral perspective (e.g., Distended fin). Bounding boxes and confidence scores indicate model detections.

## Reference

Alex, D., Cardoso, S. D., Ramos, A., & Gonçalves, D. (2023). Behavioral and endocrine responses to noninteractive live and video conspecifics in males of the Siamese fighting fish. Current Zoology, 69(5), 568. 10.1093/CZ/ZOAC078

Alton, L. A., Portugal, S. J., & White, C. R. (2012). Balancing the competing requirements of air-breathing and display behaviour during male-male interactions in Siamese fighting fish Betta splendens. 10.1016/j.cbpa.2012.11.012

Benjamini, Y., & Hochberg, Y. (1995). Controlling the False Discovery Rate: A Practical and Powerful Approach to Multiple Testing. Journal of the Royal Statistical Society: Series B (Methodological), 57(1), 289–300. 10.1111/j.2517-6161.1995.tb02031.x

da Silva, M. C., Canário, A. V. M., Hubbard, P. C., Cardoso, S. D., & Gonçalves, D. (2025). High variability in aggression and habituation to the mirror assay in ornamental Siamese fighting fish Betta splendens. Acta Ethologica, 28(3), 139–150. 10.1007/s10211-025-00469-9

Fazzari, E., Romano, D., Falchi, F., & Stefanini, C. (2025). Animal behavior analysis methods using deep learning: A survey. Expert Systems with Applications, 289, 128330. 10.1016/j.eswa.2025.128330

Friard, O., & Gamba, M. (2016). BORIS: a free, versatile open-source event-logging software for video/audio coding and live observations. Methods in Ecology and Evolution, 7(11), 1325–1330. 10.1111/2041-210X.12584

Giancardo, L., Sona, D., Huang, H., Sannino, S., Managò, F., Scheggia, D., Papaleo, F., & Murino, V. (2013). Automatic Visual Tracking and Social Behaviour Analysis with Multiple Mice. PLOS ONE, 8(9), e74557. 10.1371/journal.pone.0074557

Guidelines for the treatment of animals in behavioural research and teaching. (2012). Animal Behaviour, 83(1), 301–309. 10.1016/j.anbehav.2011.10.031

Hsieh, C.-M., Hsu, C.-H., Chen, J.-K., & Liao, L.-D. (2024). AI-powered home cage system for real-time tracking and analysis of rodent behavior. iScience, 27(11), 111223. 10.1016/j.isci.2024.111223

Hu, J., Zhao, D., Zhang, Y., Zhou, C., & Chen, W. (2021). Real-time nondestructive fish behavior detecting in mixed polyculture system using deep-learning and low-cost devices. Expert Systems with Applications, 178, 115051. 10.1016/j.eswa.2021.115051

Kendall, A., Grimes, M., & Cipolla, R. (2015). PoseNet: A Convolutional Network for Real-Time 6-DOF Camera Relocalization. 2938–2946. 10.1109/ICCV.2015.336

Krackhardt, D. (1988). Predicting with networks: Nonparametric multiple regression analysis of dyadic data. Social Networks, 10(4), 359–381. 10.1016/0378-8733(88)90004-4

Lapp, H. E., Salazar, M. G., & Champagne, F. A. (2023). Automated maternal behavior during early life in rodents (AMBER) pipeline. Scientific Reports, 13(1), 18277. 10.1038/s41598-023-45495-4

Lin, T.-Y., Maire, M., Belongie, S., Hays, J., Perona, P., Ramanan, D., Dollár, P., & Zitnick, C. L. (2014). Microsoft COCO: Common Objects in Context. In D. Fleet, T. Pajdla, B. Schiele, & T. Tuytelaars (A c. Di), Computer Vision – ECCV 2014 (pp. 740–755). Springer International Publishing. 10.1007/978-3-319-10602-1_48

Mathis, A., Mamidanna, P., Cury, K. M., Abe, T., Murthy, V. N., Mathis, M. W., & Bethge, M. (2018). DeepLabCut: Markerless pose estimation of user-defined body parts with deep learning. Nature Neuroscience, 21(9), 1281–1289. 10.1038/s41593-018-0209-y

Noldus, L. P. J. J., Spink, A. J., & Tegelenbosch, R. A. J. (2001). EthoVision: A versatile tracking system for automation of behavioral experiments. *Behavior Research Methods*, Instruments & Computers, 33(3), 398–414. 10.3758/BF03195394

Palmiotti, A., Lichak, M. R., Shih, P.-Y., Kwon, Y. M., & Bendesky, A. (2023). Genetic manipulation of betta fish. Frontiers in Genome Editing, 5. 10.3389/fgeed.2023.1167093

Pérez-Escudero, A., Vicente-Page, J., Hinz, R. C., Arganda, S., & de Polavieja, G. G. (2014). idTracker: Tracking individuals in a group by automatic identification of unmarked animals. Nature Methods, 11(7), 743–748. 10.1038/nmeth.2994

Ramos, A., Alex, D., Cardoso, S. D., & Gonçalves, D. (2021). Androgens and corticosteroids increase in response to mirror images and interacting conspecifics in males of the Siamese fighting fish Betta splendens. Hormones and Behavior, 132, 104991. 10.1016/j.yhbeh.2021.104991

Ramos, A., & Gonçalves, D. (2019). Artificial selection for male winners in the Siamese fighting fish Betta splendens correlates with high female aggression. Frontiers in Zoology, 16(1). 10.1186/s12983-019-0333-x

Ramos, A., & Gonçalves, D. (2022). Selection for winners impacts the endocrine system in the Siamese fighting fish. General and Comparative Endocrinology, 318, 113988. 10.1016/j.ygcen.2022.113988

Ribeiro, R., Cachitas, H., Mao de Ferro, C., & Teixera, L. (2019). Python video annotator [Software]. GitHub Repository. https://github.com/video-annotator/pythonvideoannotator

Simpson, M. J. A. (1968). The Display of the Siamese Fighting Fish, Betta splendens. Animal Behaviour Monographs, 1, i–73. 10.1016/s0066-1856(68)80001-9

Smith, H. M. (1945). The fresh-water fishes of Siam, or Thailand. Bulletin of the United States National Museum, (188), 1–622. 10.5479/si.03629236.188.1

Sun, G., Lyu, C., Cai, R., Yu, C., Sun, H., Schriver, K. E., Gao, L., & Li, X. (2021). DeepBhvTracking: A Novel Behavior Tracking Method for Laboratory Animals Based on Deep Learning. Frontiers in Behavioral Neuroscience, 15. 10.3389/fnbeh.2021.750894

Wei, Q., Wang, J., Zhai, G., Pang, R., Yu, H., Deng, Q., Liu, X., & Zhou, Y. (2025). PursuitNet: A deep learning model for predicting competitive pursuit-like behavior in mice. Brain Research, 1858, 149634. 10.1016/j.brainres.2025.149634

Yadav, R. S. P., Dey, P., Ansari, F., Kottat, T., Prabhu, P. P., Vasam, M., Ayyangar, S., S, S. B., Prabhu, K., Ghosh, M., & Agrawal, P. (2025). DANCE: An open-source analysis pipeline and low-cost hardware to quantify aggression and courtship in Drosophila. eLife, 14. 10.7554/eLife.105465.1

Zhang, W., Wang, H., Brandt, D. Y. C., Hu, B., Sheng, J., Wang, M., Luo, H., Li, Y., Guo, S., Sheng, B., Zeng, Q., Peng, K., Zhao, D., Jian, S., Wu, D., Wang, J., Zhao, G., Ren, J., Shi, W., … Hong, Y. (2022). The genetic architecture of phenotypic diversity in the Betta fish (Betta splendens). Science Advances, 8(38), 4955. 10.1126/SCIADV.ABM4955/SUPPL_FILE/SCIADV.ABM4955_SM.PDF

